# Social status in zebrafish modulates the behavioral response to 5-HT2C receptor agonists and antagonists

**DOI:** 10.1101/2023.04.26.538457

**Authors:** Larissa Nunes de Oliveira, Nuno Felix Paiva Alves, Marta Candeias Soares, Caio Maximino

## Abstract

The effects of previous social experiences on social behavior have been demonstrated across species both in cooperative and competitive contexts. In dominance-subordinate hierarchies, differences across social ranks have been observed in many different mechanisms. Dominance hierarchies interfere in defensive behavior, where subordinate animals present a greater defensive behavior, regarding potential threats (“anxiety-like behavior”), than dominant animals. The serotonergic system plays a key role in regulating and mediating threat responses, including 5-HT2 receptors in the types of proximal threat responses modulated by the stress of social defeat. We separated 148 adult zebrafish in pairs, and allowed to interact for five days; after that, the dominant-subordinate rank was determined, and animals were treated with a 5-HT_2C_ receptor agonist (MK-212) or antagonist (RS-102221) before being observed in the novel tank test. While MK-212 increased bottom-dwelling, erratic swimming, and freezing across all statuses, RS-102221 decreased these variables in dominants but increased them in subordinates. Moreover, the effects of MK-212 were larger in subordinates than in controls or dominants, suggesting a sensitization of the 5-HT_2C_ receptor.

## 1. Introduction

Social plasticity or competence refers to the adaptive capacity to alter behavior with changing social contexts, as well as to adapt the behavioral response according to previous social experience (Maruska et al., 2019; Oliveira, 2012; Taborsky & Oliveira, 2012). The effects of previous social experiences on social behavior have been demonstrated across species both in cooperative and competitive contexts; for example, social fish are able to adapt their aggression levels based on previous experience as “winner” or “loser” in agonistic encounters (Oliveira et al., 2011, 2016), as well as from gathering information on opponents while observing fights (Doutrelant et al., 1998; Oliveira et al., 1998, 2016).

Conversely, reciprocity-based adjustments in cooperative behavior are also observed, with animals cooperating more frequently with individuals which cooperated with them in the past (Dugatkin, 1998; Pimentel et al., 2019, 2021). Other examples exist in which experiences with specific social environments, lead to changes in non-social behavior. For example, changes in sensory responsiveness were observed in *Astalotilapia burtoni* after the establishment of dominance hierarchies (Maruska & Fernald, 2010; Nikonov et al., 2017; Nikonov & Maruska, 2019). Dominance hierarchies also alter defensive behavior, with subordinate animals typically showing higher levels of responses to potential threats (“anxiety-like behavior”) than dominant animals (Blanchard et al., 1993; Bozi et al., 2021). Social plasticity thus allows animals to flexibly change their behavior in relation to social demands, which suggests important consequences for fitness (Taborsky & Oliveira, 2012).

The proximate mechanisms of social plasticity have been studied in diverse vertebrate taxa; studies in fish are of special interest because this diverse group occupies a wide range of ecological and ecosocial niches, with great variety of social arrangements (Maruska et al., 2019). In general, a “core” mechanism for social plasticity across fish species is the initiation of behavioral shifts through the modulation of a “social decision-making network” by monoamines, neuropeptides, and steroid hormones (Maruska et al., 2019; Oliveira, 2012); the consolidation of these rapid shifts is mediated mainly by neurogenomic adjustments and regulation of the expression of transcription factors, cell cycle regulators, and plasticity products (Cardoso et al., 2015; Maruska et al., 2019).

The social decision-making network (SDMN) is a neural survival circuit involved in both reward and sociality that appears to be largely conserved across vertebrate taxa (O’Connell & Hofmann, 2011, 2012). The minimum circuit of the SDMN involves both the social behavior network and the mesolimbic reward system, with an overlap on brain regions involved in the regulation and integration of responses to salient social and non-social stimuli. In mammals, the SDNM involves the lateral septum, extended medial amygdala and bed nucleus of the stria terminalis, preoptic area/paraventricular nucleus (POA/PVN), anterior hypothalamus, ventromedial hypothalamus, and periaqueductal gray area, as well as six areas of the mesolimbic reward system – the striatum, nucleus accumbens, ventral pallidum, basolateral amygdala, hippocampus, and ventral tegmental area (O’Connell & Hofmann, 2011, 2012). In teleost fish, nodes of this network have been identified, although only partial homologies have been established for many of the regions involved. These include the dorsomedial telencephalon (Dm), dorsolateral telecenphalon (Dl), ventral subpallium (Vv), dorsal subpallium (Vd), central subpallium (Vc), supra- and post-commissural subpallium (Vs and Vp), preoptic area (POA), ventral tuberal nucleus (VTn), anterior tuberal nucleus (ATn), and central gray (GC)(O’Connell & Hofmann, 2011). Interestingly, some nodes of the social decision-making network (Dm, Vv, Vs, Vp, POA, and GC) are also part of an “aversive brain system” involved in mediating responses to threatening stimuli (do Carmo Silva et al., 2018), which could represent a mechanism through which stress and sociality interact (Soares et al., 2018).

In dominance-subordinate hierarchies – the case that will be studied in the present work –, differences across social ranks have been observed in many different mechanisms. In the brains of dominant zebrafish (*Danio rerio*), the mRNA levels of *tyrosine hydroxylase 2, hypocretin/orexin* and *histidine decarboxylase* were increased in relation to subordinates, which had increased levels of *arginine vasotocin* (Pavlidis et al., 2011). Conversely, subordinate animals had higher mRNA levels of genes involved in serotonergic signalling, including *htr1aa* (which encodes one isoform of the 5-HT_1A_ receptor), *htr1b* (which encodes the 5-HT_1B_ receptor), *htr2b* (which encodes the 5-HT_2B_ receptor), and *slc6a4b* (which encodes one isoform of the serotonin transporter that is expressed mainly in serotonergic neurons of the hypothalamus; Norton et al., 2008), while dominant animals showed higher mRNA levels of *slc6a4a* (which encodes one isoform of the serotonin transporter that is expressed mainly in serotonergic neurons of the raphe and the pretectal cluster; Norton et al., 2008).

In salmonids, while the initial agonistic encounters lead to a rapid activation of the brain serotonergic system in both dominant and subordinate individuals, in absence of signs of altered defensive behavior, as soon as the hierarchy is established serotonergic activity returns to baseline levels in dominant fish, but not subordinate animals (Øverli et al., 1999). These differences are examples of social plasticity related to the experience of winning or losing an encounter; indeed, in zebrafish serotonergic activity is significantly higher in the telencephalon of winners and in the optic tectum of losers, while no effects were observed in animals fighting against a mirror image (Teles et al., 2013). In agreement with that observation, Dahlbom et al. (2012) showed that subordinate zebrafish had higher serotonergic activity in the hindbrain than dominants at the fifth day of interaction, when hierarchies are clearly established.

Given the importance of the serotonergic system in organizing responses to threat across vertebrates (Graeff et al., 1997; Herculano & Maximino, 2014; Krause et al., 2017), it is likely that these effects of social rank mediate social plasticity of defensive behavior. In zebrafish, a dual role for the serotonergic system has been shown for defensive behavior, with serotonin increasing responses to potential threats and decreasing responses to distal or proximal threats (Lima-Maximino et al., 2020; Maximino et al., 2013). Both 5-HT_1A_ receptor agonists and antagonists have been shown to decrease responses to potential threats in non-stressed zebrafish (Araujo et al., 2012; Maximino et al., 2013), an effect that was like that of 5-HT_1B_ receptor antagonists or inverse agonists (Maximino et al., 2013; Nowicki et al., 2014). 5-HT_2C_ receptor agonists were shown to reduce responses to distal threat (conspecific alarm substance), while 5-HT_2C_ receptor antagonists decreased responses to a potential threat (novel tank after exposure to the alarm substance) (do Carmo Silva et al., 2021). Serotonin also decreases mirror-elicited aggression in zebrafish (Theodoridi et al., 2017), which could indicate a role for the serotonergic tone in regulating the suppression of aggressive behavior observed in subordinate individuals.

These latter effects are partially mediated by the 5-HT_2C_ receptor, given that RS-102221, a 5-HT_2C_ receptor antagonist, decreased time in the display zone in the mirror-induced aggressive display test, but increased display duration (Moura et al., 2023). Indeed, a role of this receptor in social behavior has been shown in zebrafish, with activation of the receptor increasing social preference and decreasing cooperation, while blocking the receptor inhibits social investigation but increases preference for social novelty (Moura et al., 2023).

Backström and Winberg (2017) describe the behavioral phenotype of subordinate individuals as “behavioral inhibition", including appetite suppression, reduced aggression, and decreased reproductive behavior. The term also suggests behavioral tendencies associated with defenses against proximal threat, including “anxiety-like behavior” (McNaughton & Corr, 2004). Indeed, we have previously shown that subordinate zebrafish show increased anxiety-like behavior in the novel tank test in relation to both controls and dominant animals, an effect that was sex-independent (Bozi et al., 2021).

Given the role of the serotonergic system in regulating and mediating responses to threat, and the recently uncovered role of 5-HT_2_ receptors in the types of responses to proximal threat which are modulated by social defeat stress (Silva et al., 2021) as well as its complex roles in sociality (Ponzoni et al., 2016; Moura et al., 2023), the aim of the present work was to elucidate the participation of 5-HT_2C_ receptors on social rank-modulated differences in responses to a potential threat. In order to do that, animals were separated in pairs, and allowed to interact for five days; after that, the dominant-subordinate rank was determined, and animals were treated with a 5-HT_2C_ receptor agonist (MK-212) or antagonist (RS-102221) before being observed in the novel tank test, an widely-used test to assay responses to potential threat/anxiety-like behavior (Bencan et al., 2009; Cachat et al., 2011; Levin et al., 2007). We found that, while MK-212 increased bottom-dwelling, erratic swimming, and freezing across all statuses, RS-102221 decreased these variables in dominants but increased them in subordinates. Moreover, the effects of MK-212 were larger in subordinates than in controls or dominants, suggesting a sensitization of the 5-HT_2C_ receptor.

## 2. Materials and methods

### 2.1. Animals and housing

In total, 148 adult zebrafish were used for this experiment. Animals were obtained from a commercial breeder licensed for aquaculture and kept in laboratory for 4 weeks for acclimatization before the beginning of the experiments. During this acclimatization, the individuals were kept in mixed-sex 40 L tanks, with a maximum density of 25 fish per tank, and in a 14h light/10h dark photoperiod. The water present in the tanks was kept at room temperature (28 ºC), with a pH of 7.0-8.0, and water parameters were routinely monitored and kept at recommended levels (Lawrence, 2007). All experiments were carried out with consideration for fish welfare and approved by UEPA’s IACUC under protocol 06/18.

### 2.2. Drugs and treatments

The individuals underwent one of the following treatments: Cortland’s salt solution, DMSO, MK-212 (2 mg/kg), or RS-102221 (2 mg/kg). The 5-HT_2C_ receptor agonist MK-212 was bought from Sigma-Aldrich (St Louis, USA) and dissolved in Cortland’s salt solution (NaCl 124.1 mM, KCl 5.1 mM, Na2HPO4 2.9 mM, MgSO4 1.9mM, CaCl2 1.4 mM, NaHCO3 11.9 mM, Polyvinylpyrrolidone 4%, 1,000 USP units Heparin) (Wolf, 1963). The 5-HT_2C_ receptor antagonist RS-102221 was bought from Sigma-Aldrich (St Louis, USA) and dissolved in DMSO. Cortland’s solution was used as a control for MK-212 and DMSO as a control for RS-102221, in accordance to their solubility. The treatments were administrated intraperitoneally and were randomly distributed in order to assure that, in each round of dyadic encounters, there were always individuals that received each one of the 4 treatments. Animals were injected with one of the drugs or either of the vehicles immediately after recording of the last contest, and left to rest for 30 min. for the onset of the drug effect before being subjected to the novel tank test (Fig.1).

**Figure 1.**
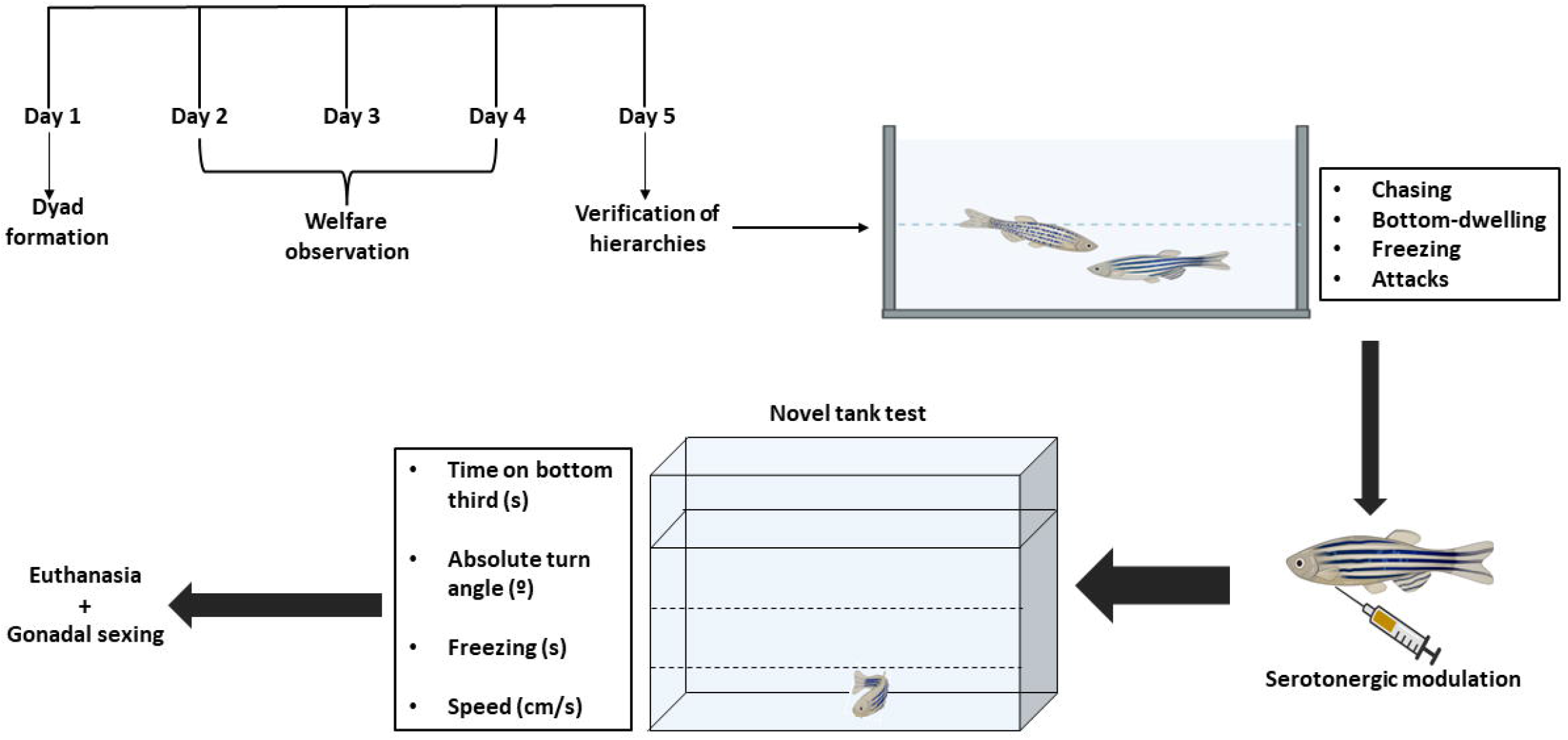
Representation of the experimental design. Dyads were allowed to be established on the experimental tanks on Day 1 and verification of hierarchy formation was done on Day 5 (the welfare of the individuals were monitored in between those days). The individuals were then moved into a novel tank test and observed for anxiety-like behavior. Following the test, gonadal sexing took place. See “Material and Methods” for further information. Image adapted from Bozi et al. (2021).

### 2.3. Dyadic encounters and establishment of hierarchy

On day 0, immediately prior to dusk, 20 individuals were randomly moved in pairs to 10 experimental tanks (20 cm × 14.5 cm × 12.5 cm) and left overnight to acclimatize with the tank, recover from stress and return to normal behavior. They were kept in these tanks for the next 5 straight days and allowed to establish dominance hierarchies. Throughout the 5 days, the welfare of the individuals was observed and maintained. This time period has been shown in previous experiments to be sufficient for the establishment of hierarchies in this species (Bozi et al., 2021; Filby et al., 2010; Paull et al., 2010). On the morning of the 5th day, all experimental tanks were video-recorded for 5 minutes and X-Plo-Rat (Tejada, Chaim, & Morato, 2017) was used to assess different behaviors, following the work described in Pavlidis et al. (2011) to determine dominant and subordinate status.

The assessed behaviors were: 1) Chasing - the number of times one of the individuals swims directly toward the other individual and pursues it during the recorded 5 minutes (Paull et al., 2010); 2) Bottom-dwelling defined as the time the fish spent on the bottom third of the experimental tank (Bozi et al., 2021); 3) Freezing, defined as the time spent without moving in any direction (Blaser & Gerlai, 2006); and 4) Attacks – the number of agonistic events, with or without association to biting (Paull et al., 2010) (Fig.1). The control fish (n=48) were subjected to the treatments but were not subjected to the dyadic encounters, having been transferred directly to the novel tank for the next phase of the experiments.

After the video analysis, the individual that exhibited the highest frequency of chasing events and attacks and the lowest time spent freezing and/or bottom-dwelling was considered the dominant one from the pair, and the individual that showed the inverse frequencies was considered to be the subordinate. This procedure was then repeated for another 8 rounds, signifying a total of 148 individuals used in the experiment.

### 2.4. Novel Tank Test (NTT)

After ascribing dominant or subordinate status to the experimented individuals and injections, all animals (including the ones used as control) were carried out to the following phase of the procedures, the Novel Tank Test, used to assess anxiety-like behavior (Bencan et al., 2009). Each animal was individually transported to a new aquarium (25 cm × 24 cm × 20 cm) filled with 5 L of system water and allowed to freely explore the apparatus for 6 minutes. Their behavior during this interval was recorded using a video camera positioned directly in front of the tank. The video recordings were filed and analyzed using TheRealFishTracker, an automated video tracking software (available from http://www.dgp.toronto.edu/~mccrae/projects/FishTracker/). The assessed behavioral variables were the time spent on the bottom third of the tank (measured in seconds); erratic swimming (absolute turn angle, measured in degrees (Bozi et al., 2021; Tran et al., 2017)); total amount of time spent freezing (measured in seconds), which was considered to happen whenever the individual would swim at a speed lower than 0.5 cm/s; and the average swimming speed (measured in cm/s) (Fig.1). The time spent on the bottom third of the tank was the primary outcome, while the remaining variables were secondary. Speed was used as an outcome to assess any potential psychomotor effects of the treatments.

### 2.5. Euthanasia and gonadal sexing

Following the experiment, the sex of the individuals was confirmed via observation of the individuals’ gonadal tissue under a light microscope (Leica CTR4000). The animals were sacrificed by immersing them in ice-cold water, followed by spinal transection (Matthews & Varga, 2012).

The individuals were then fixed in Bouin’s fixative for 24h. To be better fixed, an incision was made on the ventral side of the individual. The fixed tissues were included in meth-acrylate glycol and sections were cut at 5 μm slices, which were collected onto glass slides. Following these steps, the slices were stained with hematoxylin-eosin to be observed under a microscope (Bozi et al., 2021). The oocytes were classified following the guidelines and descriptions presented in Selman et al. (1993) (Fig. 1).

### 2.6. Statistical Analysis

The information regarding sex, treatment, status and measurements of the dependent variables (time spent in bottom third of the tank, absolute turn angle, freezing and speed) were compiled onto a table. The table was then imported to R 4.1.2 using Rstudio. The data were first plotted to observe any qualitative differences between males and females in the different groups, following Joel & Fausto-Sterling (2016). The rationale for visual inspection is that there is evidence suggesting that differences seen between male and female brains may be insufficient and misleading for establishing relationships between sex and brain (Joel & Fausto-Sterling, 2016). Furthermore, these differences (both in humans and animals used in scientific research) may arise from one heterogeneous group rather than two separate ones. As such, it is suggested to observe for any possible interactions between sex and treatment and, if no interactions are suggested, the sex category should be omitted from the results’ analysis (Joel & Fausto-Sterling, 2016). The preliminary plots showed no major differences between sexes in any group (Supplemental Figure 1) and, as such, the variable “Sex” was not included in the subsequent analyses.

Following that step, data were analyzed using two-way ANOVAs on trimmed means (Wilcox, 2012), since assumptions of normality and homoscedasticity were violated for most endpoints. These analyses were made on R 4.1.2 using the package WRS2 (v. 1.4; https://r-forge.r-project.org/projects/psychor/).

## 3. Results

Both social status (Q = 467.63, p < 0.001) and treatment (Q = 354.17, p < 0.001) had significant effects on the time spent in the bottom third of the tank (Figure 2A). A significant interaction was also found for this variable (Q = 162.35, p < 0.001). The post-hoc tests showed that control and dominant individuals behave similarly when comparing Cortland’s solution vs DMSO (p=0.176), and the same was found when comparing control and subordinate individuals (p=0.648). However, the difference in the time spent at the bottom significantly differed between dominant individuals and subordinates (p=0.025). MK-212 increased bottom-dwelling across all treatments (p < 0.01), with a smaller effect in subordinate animals (d = -1.22, 95%CI[-2.04, -0.4]) than in both controls (d = -3.2, 95%CI[-4.09, -2.31) and dominant animals (d = -2.89, 95%CI[-3.74, -2.04]). Treatment with RS-102221 decreased the time spent in the bottom third in controls (p=0.012; d =2.51, 95%CI[1.65, 3.37]) and dominant animals (p=0.006; d =1.61, 95%CI[0.81, 2.41). In subordinates, the 5-HT_2C_ antagonist synergistically increased the time spent in the bottom third in comparison to the respective control-treated individuals (p = 0.004; d =-3.04, 95%CI[-3.93, 02.15]).

**Figure 2.**
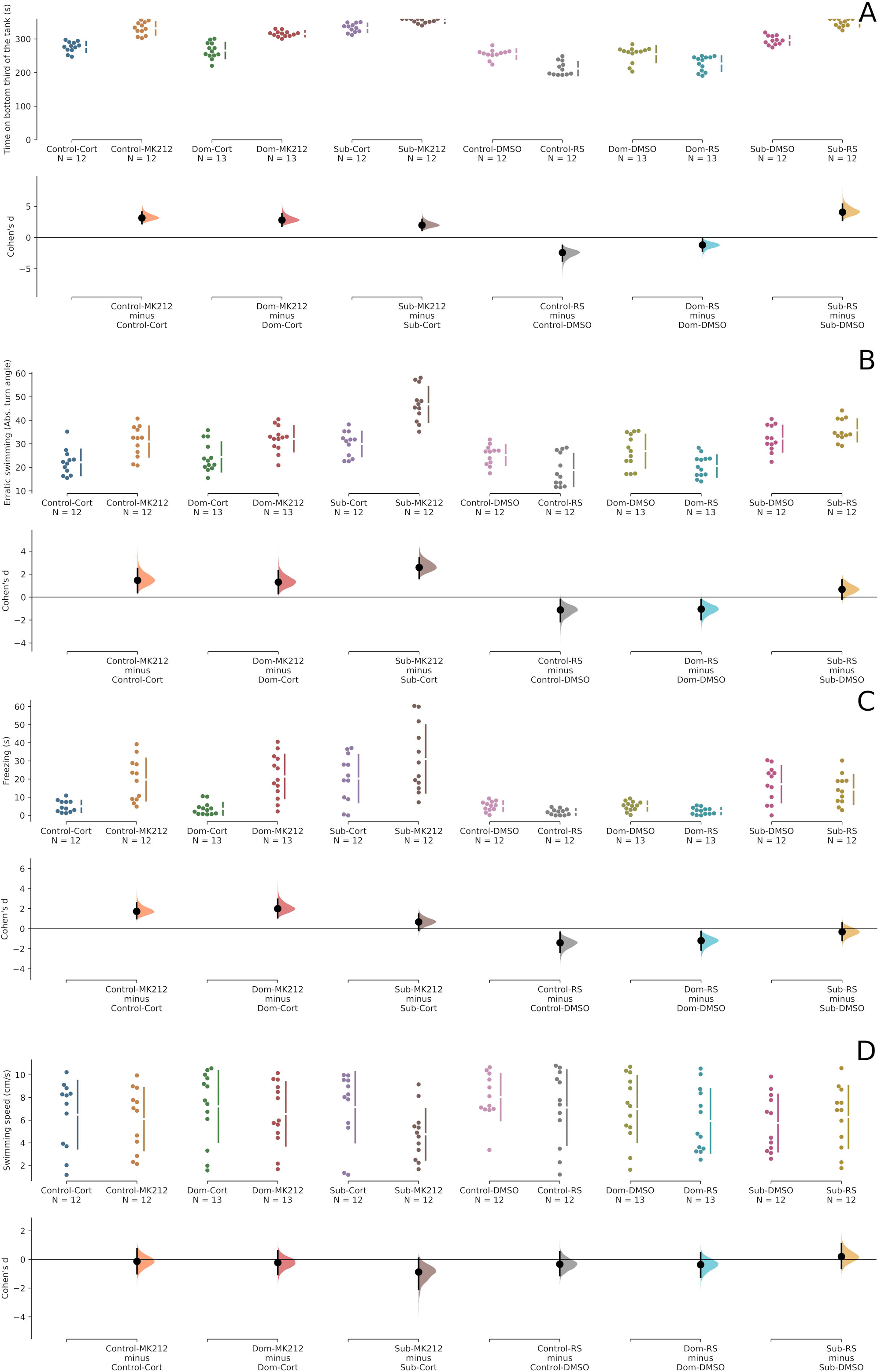
Effects of social status and drug treatments on (A) Geotaxis, (B) Erratic swimming, (C) Freezing, and (D) Swimming speed. Cohen’s *d*s for 6 comparisons are shown in the Cumming estimation plots. The raw data is plotted on the upper axes; each mean difference is plotted on the lower axes as a bootstrap sampling distribution. Mean differences are depicted as dots; 95% confidence intervals are indicated by the ends of the vertical error bars.

Regarding erratic swimming, both social status (Q = 73.13, p < 0.001) and drug treatment (Q = 58.53, p < 0.001) had significant effects (Figure 2B). However, no significant interaction effects were found (Q = 13.31, p = 0.09). The post-hoc tests revealed significant differences between social status (control vs. subordinates: p<0.001, d = -2.01, 95%CI[-2.48, -1.54]; dominants vs. subordinates: p<0.001, d =-1.72, 95%CI[-2.17, -1.27]; controls vs. dominants: p=0.226, d = - 0.29, 95%CI[-0.69, 0.11]) in erratic swimming. Subordinate individuals displayed more erratic swimming compared to control and dominant individuals, and MK-212 increased erratic swimming across all statuses (p < 0.01, d = -1.88, 95%CI[-2.39, -1.37]). RS-102221 did not affect erratic swimming.

Effects of both social status (Q = 27.19, p < 0.001) and drug treatment (Q = 34.71, p < 0.001) were found to be significant regarding freezing (Figure 2C), but no interaction was found (Q = 4.31, p = 0.689). The post-hoc tests revealed significant differences between individuals of different social statuses, with subordinates freezing more than both controls and dominants (controls vs. subordinates: p<0.001, d = -1.39, 95%CI[-1.83, -0.95]; dominants vs. subordinates: p<0.001, d = -1.38, 95%CI[-1.81, -0.95]; controls vs. dominants: p = 0.999, d = -0.01, 95%CI[-0.4, 0.39]). Across social statuses, MK-212 increased freezing (p < 0.001, d = -1.57, 95%CI[-2.07, - 1.07]), but no effect was found for RS-102221 (p = 0.6, d = 0.3, 95%CI[-0.16, 0.76]).

Finally, no main effects of status (Q = 3.11, p = 0.225) nor drug treatment (Q = 5.30, p = 0.177) were found for swimming speed (Figure 2D). An interaction effect was also absent (Q = 8.16, p = 0.303).

## 4. Discussion

In this study we aimed to find the involvement of 5-HT_2C_ receptors on social rank-modulated differences regarding anxiety-like responses to a potential threat. For that, individuals were set up in pairs and allowed to establish dominance hierarchies, which were confirmed through behavioral analysis in the 5th day. After ascribing dominant or subordinate status, all experimented individuals were injected and then tested using the Novel Tank test. MK-212 was found to increase bottom-dwelling, erratic swimming, and freezing behavior for dominants and subordinates, while RS-102221 decreased these variables solely in dominants but otherwise increased them in subordinates. Moreover, the effects of MK-212 were also more significant in subordinates than in dominants and controls.

Similarly to evidence by Bozi and colleagues (2021), we found that becoming subordinate lead to more geotaxis, erratic swimming, and freezing behavior, as it is considered a psychosocial stressor. Interestingly, these effects are also not likely due to changes in activity, as swimming speed was not affected by any treatment. Other stressor types (such as restraint stress or alarm substances) have also been shown to result in increased geotaxis, freezing, and erratic swimming in stressed individuals (Ghisleni et al., 2012; Lima-Maximino et al., 2020; Silva et al., 2021). Increased freezing behavior in subordinates can also been associated with a reactive or passive coping style, which is typical of subordinate individuals. This association seems evolutionarily conserved, as subordinate rats in diverse social stress paradigms showcase passive defensive strategies highlighted by freezing for most of the time (Motta et al., 2009; Nyuyki, Beiderbeck, Lukas, Neumann, & Reber, 2012).

In this study, no major differences were found between sexes after preliminary, qualitative observations in all behavioral variables studied. Similar results were observed in other studies in zebrafish, in which differences between sexes were absent in regard to the effect of social status on anxiety (Bozi et al., 2021). Moreover, the influence of sex in the behavior of this species has not been consistent. For example, Tea et al. (2019) showed that male subordinate zebrafish were shown to have higher cortisol levels and reduced cell proliferation in the forebrain, which was not seen in female subordinates. However, the authors did not focus on anxiety-like behavior (Tea et al, 200)). In rodents, the induction of social stress by forming hierarchies is not always visible in females, given that laboratory female rats are usually less aggressive and less likely to establish hierarchies (Tamashiro et al., 2005); nevertheless, that does not appear to be the case in the present work, as both males and females were able to establish hierarchies – including same- and different-sex hierarchies. Again, similar effects were observed in previous work by our group (Bozi et al., 2021). These overall discrepancies between studies should warrant further analyses, as it has also been suggested recently in Genario et al. (2020).

Regarding our pharmacological manipulations, differences were also found between control vehicles in geotaxis. In fact, the average time spent in the bottom third by DMSO-treated individuals was lower than the average time by Cortland’s solution-treated individuals for all social status. Given that Cortland’s solution is a saline solution, this could possibly be attributable to a relatively mild osmotic shock created by it that doesn’t impact much dominant individuals, but, in subordinates, has an additive impact in this variable to the effect caused by their social rank. However, this should mostly be associated to the effect of DMSO, as this solution has shown to induce reductions of anxiety-like behaviors in the novel tank test (Nowicki et al., 2014), as well as in the light/dark plus maze test (Sackerman et al., 2010).

Results on geotaxis, the main endpoint of the novel tank test, revealed significant differences between dominant and subordinate individuals, with subordinates spending more time spent in the bottom third of the tank, regardless of treatment, compared to dominant individuals. This is an expected result, as it replicates previous works. For example, in Bozi et al. (2021), subordinates showed higher geotaxis than dominants. Interestingly, 5-HT_2C_ receptor activation via MK-212 treatment resulted in increased geotaxis not only in subordinate individuals, but also in control and dominant individuals. This is consistent with a general anxiogenic effect of MK-212 independent of social status. Likewise, MK-212 increased erratic swimming and freezing independently of social status. MK-212 was shown increase anxiety-like behavior in rats in the elevated plus-maze (Alves et al., 2004; Cruz et al., 2005). This drug also shows anxiogenic effects in humans (Lowy & Meltzer, 1988). In zebrafish, MK-212 blocked the effects of conspecific alarm response on freezing and geotaxis (Carmo Silva et al., 2021), but no effects were observed on post-exposure behavior, a model used to assess responses to potential threat / anxiety-like behavior.

When effect sizes were compared, the effects of MK-212 on geotaxis were shown to be larger in dominants than subordinates. This could be explained either by a ceiling effect on subordinate individuals, or by an important interaction between social rank and the treatment. The first hypothesis cannot be discarded since, in vehicle-treated subordinates, the mean geotaxis is close to 360 seconds, which is the upper limit for the measurements in this variable. As such, the rise in geotaxis caused by the treatment with MK-212 in subordinates could not be very large (ceiling effect). If ceiling effects are not the main driver of these differences, these may also arise through a desensitization of 5-HT_2C_ receptors in subordinate individuals. For instance, 5-HT_2_ receptors appear to decrease their responsiveness upon chronic agonist stimulation by decreasing receptor density (van Oekelen et al., 2003). Thus, if the experience of social stress increases serotonergic activity in the hindbrain (Dahlbom et al., 2012), it is possible that the 5-HT_2C_ receptors are desensitized by the fifth day of interaction, reducing the effect of MK-212 in subordinates.

In controls and dominant individuals, the 5-HT_2C_ receptor RS-102221 decreased geotaxis, erratic swimming, and freezing, which is consistent with the anxiolytic properties of this drug (Bell et al., 2014; Kuznetsova, Amstislavskaya, Shefer, et al., 2006; Scarpelli et al., 2008; Warhaftig et al., 2021). In subordinates, on the other hand, RS-102221 synergistically increased these variables. If the reduced effects of MK-212 on geotaxis in subordinate individuals are due to receptor desensitization or down-regulation associated with the elevated serotonergic tone in the hindbrain, we should expect the effects of RS-102221to also be reduced in subordinates. Thus, it is likely that the differential effects of MK-212 and RS-102221 on anxiety-like behavior in subordinate zebrafish reflect the behavioral constraints associated with the contingencies of hierarchy formation. Status-dependent differences in the effects of drugs have been previously reported in mammals (e.g., Raleigh et al., 1985; Winslow & Insel, 1991); for example, Raleigh et al. (1985) found that drugs that increase serotonergic neurotransmission (fluoxetine, tryptophan, or quipazine) were more potent in changing the behavior of dominant than subordinate vervet monkeys (*Chlorocebus pygerythrus*). These effects are reminiscent of the reduced effects of MK-212 in subordinate zebrafish found in the present experiments. The differences in effects between MK-212 and RS-102221 could also be related to differences in receptor sensitivity and/or expression across brain regions; while the expression of 5-HT_2C_-like receptors in the brains of adult zebrafish are still unknown, in larvae several clusters are found, including in the hindbrain (where serotonergic activity is higher in subordinate animals; Dahlbom et al., 2012), but also in forebrain regions such as the preoptic nucleus, ventral thalamus, and posterior tuberculum (Schneider et al., 2012).

In summary, the present results found that the behavioral effects of MK-212, a 5-HT_2C_ agonist, and RS-102221, a 5-HT_2C_ antagonist, depend on social status in adult zebrafish, with MK-212 being less effective in increasing anxiety-like behavior in subordinate animals, and RS-102221 synergistically increasing anxiety-like behavior instead of decreasing it in subordinate animals. These status-related behavioral effects of 5-HT_2C_ receptor-targeting drugs may reflect differences in receptor expression in the brains of dominant and subordinate individuals, or may otherwise depend on the behavioral differences associated with social organization. Overall, these results show a role for 5-HT_2C_ receptors in social plasticity in a model vertebrate.

## Funding

MCS is supported by Portuguese National Funds through the Fundação para a Ciência e Tecnologia – FCT/Portugal (CEEC Individual – 2021.01458.CEECIND). CM is supported by Conselho Nacional de Desenvolvimento Científico e Tecnológico – CNPq/Brazil (productivity scholarship #302998/2019-5).

## Conflicts of interest statement

Authors declare no conflict of interest

## Figure captions

Figure S1 – Average group values separated by social status, drug treatment, and sex.

## Notes

https://github.com/lanec-unifesspa/5HT2C-sociality/tree/main/hierarchy

### Competing Interest Statement

The authors have declared no competing interest.

