## Supplementary figures and images for "Social status in zebrafish modulates the behavioral response to 5-HT2C receptor agonists and antagonists"

### Supplemental Figure 1

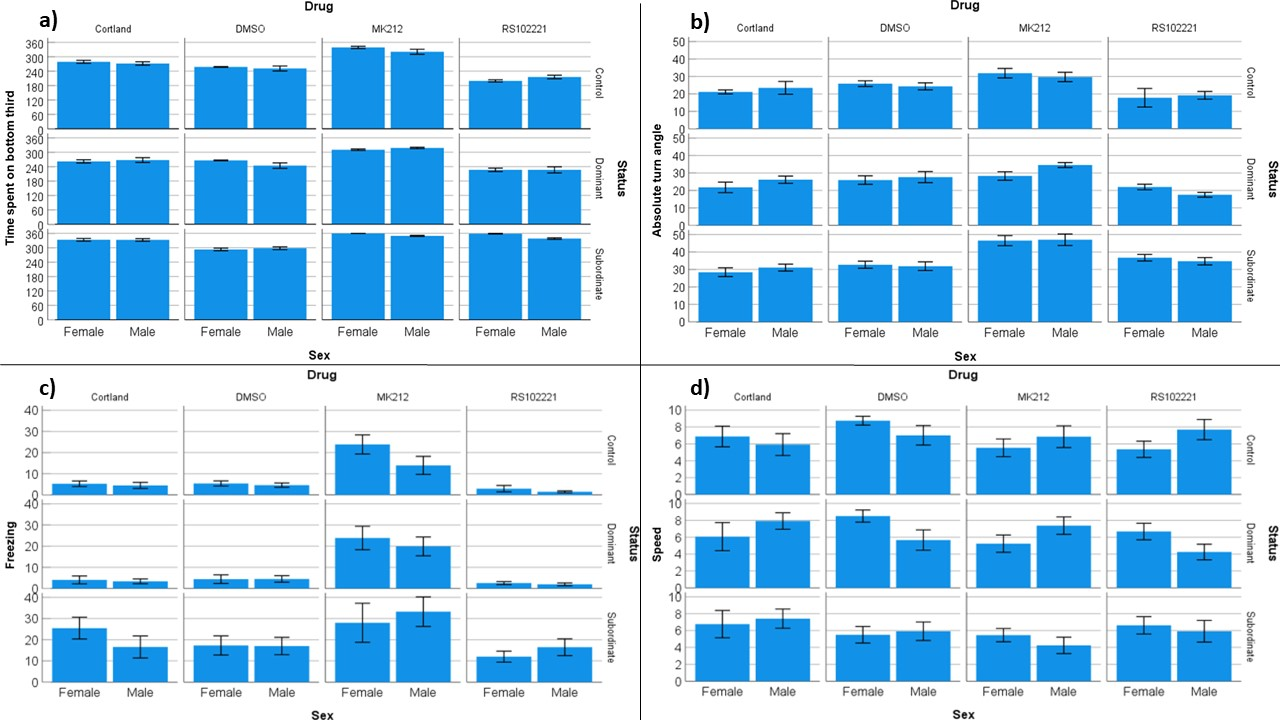
